# Evaluating Cochlear Implant Stimulation Strategies Through Wide-field Calcium Imaging of the Auditory Cortex

**DOI:** 10.1101/2024.02.05.577161

**Authors:** Bruno Castellaro, Tony Ka Wing Yip, Fei Peng, Zeeshan Muhammad, Shiyi Fang, Israel Nelken, Jan Schnupp

## Abstract

Cochlear Implants (CI) are an effective neuroprosthesis for humans with profound hearing loss, enabling deaf adults to have phone calls without lipreading and babies to have successful language development. However, CIs have significant limitations in complex hearing situations, motivating the need for further research, including studies in animal models. Here, we demonstrate the usefulness of wide field Ca++ imaging in assessing different CI stimulation strategies. One major challenge in electrophysiology in CI animals lies in excluding the CI electric artifacts from the recording, since they are orders of magnitude larger than the amplitude of action potentials. Also, electrophysiology can rarely sample large areas of neuropil at high spatial resolution. To circumvent these problems, we have set up an imaging system allowing us to monitor neural activity in the auditory cortex (AC) of CI supplied rats using the Ca++ sensitive dye OGB. Here we describe an initial experiment with this setup, in which we recorded cortical responses to 4 different stimulation patterns which were delivered across 3 CI channels to the contralateral ear. We then investigated two parameters that have been shown to affect intelligibility in CI users: pulse rate and relative pulse timing across CI channels. While pulse rate had only a very modest effect on the discriminability of the neural responses, the stimulation mode had a major effect, with simultaneous pulse timing, perhaps surprisingly, allowing much better pattern discrimination than interleaved sampling. The result suggests that allowing collisions of pulses on neighboring channels may not always be detrimental, at least if partial overlaps of pulses, in which anodic and cathodic pulse phases might cancel, are avoided.

## Introduction

Cochlear Implants (CIs) are an effective neuroprosthesis for humans with profound hearing loss, enabling many deaf adults to have phone conversations without lipreading and enabling deaf babies to successfully acquire spoken language. However, CI hearing underperforms relative to natural hearing in complex hearing situations such as music appreciation, sound localization, speech in noise detection, or perception of tonal languages (Riley P. et al. 2018; Ludwig A. et al. 2021; Choi et al. 2022). It remains an important open research question whether improved electrical stimulation strategies might permit significant improvements in the quality of auditory perception afforded by CIs. The current state of the art, summarized in a recent review by Carlyon R. and Goehring (2021), remains largely based on the Continuous Interleaved Sampling (CIS) strategy invented by (Wilson et al. 1991), or a number of variants of that strategy, such as “n-of-m”. This strategy has shown significant successes, but also suffers from significant shortcomings. Even devices that seek to provide temporal fine structure information in pulse timing, such as the FS4 strategy by MedEl, do so on only a minority of channels, with most channels running a type of CIS. Consequently, there haven’t been significant departures from CIS-style processing strategies for several decades. This may be due, in part, to the challenge of testing new strategies in behavioral experiments on humans once the old strategies are firmly established. Evaluating new strategies often requires testing them on patients who are accustomed to the established strategy. However, CI users are known to require practice periods extending for several months to adapt to the stimulation patterns provided by their devices (Svirsky M. et al. 2001), and it is not realistic to expect patients to devote as much time and effort to adapt to a novel experimental strategy as they have already invested in learning to listening through the established strategies built into their clinical devices. This makes psychoacoustic comparisons of novel strategies inherently biased, as unfamiliar input will have to compete against stimulation patterns for which the participants’ auditory pathways have become optimized over many months of practice.

Animal experiments can therefore play an important role, at least in the initial evaluation of different strategies, as they provide a unique opportunity to test different stimulation strategies in an unbiased manner and in a highly controlled setting. However, animal experiments too have their difficulties. Behavioral testing of experimental animals with CIs is technically difficult and time consuming, so although animal behavior studies have led to perhaps surprising and interesting insights (Miller et al. 1995; Rosskothen-Kuhl et al. 2021; Buck A. et al. 2023), the use of this approach remains relatively rare. Electrophysiological recordings from neurons of the auditory pathway of experimental animals are somewhat easier to achieve and more commonly performed, but despite recent technical advances, this approach remains plagued by difficulties in removing the electrical stimulation artifacts generated by the CI stimuli, particularly when stimulation parameters are to be compared in the context of rich, multichannel stimulation paradigms which generate dense and complex electrical artifacts. While sophisticated techniques for removing electrical artifacts from physiological recordings exist (O’Shea and Shenoy 2018; Najafabadi M. et al. 2020), their use becomes more and more difficult as cochlear stimulation patterns increase in complexity, with more channels and faster pulse rates, and they cannot always give an entirely complete and uncontaminated picture of the underlying neural activity.

In this paper we wanted to evaluate optical methods for monitoring neural activity as a potentially useful alternative, or complementary, approach for investigating the effectiveness of different parameters of intracochlear electrical stimulation. A big advantage of such imaging methods is that they are completely unaffected by electrical stimulation artifacts, providing an uncontaminated view of neural responses to different CI stimulation patterns, and they can sample large areas of auditory cortex simultaneously, making it unlikely that interesting neural responses are missed because the recording probe is not optimally placed. Optical recording methods are likely to be particularly valuable for investigating responses to rich, multichannel stimulation patterns delivered at high pulse rates where artifact removal from electrical recordings becomes challenging. While the use of optical imaging methods to observe activity in the auditory cortex is very well established, it has not, to our knowledge, been previously used to evaluate different CI stimulation strategies.

Here we wanted to assess the advantages and disadvantages of using a relatively straight-forward optical recording technique, the monitoring of cortical population activity using wide-field imaging and the Ca++ dependent indicator dye Oregon Bapta Green (OGB). To do so, we designed a study which rests on the observation that differences in cortical population responses correlate with perceptual discriminability of important complex sounds such as speech syllables (Engineer C. et al. 2008; Shetake J. et al. 2011; Centanni et al. 2014). In the context of CI research, we therefore thought to use optical methods to observe patterns of cortical activation evoked by multichannel cochlear implant stimulation, and to quantify the extent to which different stimulation patterns evoke reliably different (and therefore discriminable) response patterns. The inherent assumption here is that the size of the differences in cortical activation patterns evoked by two different CI stimuli is likely to be predictive of how easy these stimulation patterns should be to discriminate perceptually. Under this assumption, it is then of interest to investigate how changes in stimulation parameters affect the discriminability of evoked cortical activity patterns. To put these aims into practice, we designed a set of four different multichannel CI patterns, which were delivered with two different stimulation strategies (Continuous Interleaved Sampling, CIS, and Simultaneous Sampling, SS) and two different pulse rates (300 and 1800 pps).

## Methods

### Animal subjects

Six young adult female Wistar rats were used to perform in vivo wide-field calcium imaging with CI stimulus. The mean age at the time of calcium imaging was 12 ± 3 weeks and the averaged weight was 270 ± 30 grams. All rats were purchased from the Chinese University of Hong Kong. All experimental procedures were performed under license from the Hong Kong Government Department of Health and approved by the Animal Research Ethics Sub-Committee of the City University of Hong Kong. Open field broad-band click Auditory Brainstem responses (ABRs) were recorded to verify that the animals had normal hearing prior to the experiment. We considered a rat as normal hearing if we could see ABRs at 20 dB SPL or below.

### Anesthesia

A previous study had reported that repeated Ketamine/Xylazine injections may inhibit neural responses detected with the widefield Ca++ imaging technique (Thrane et al. 2012), so we used i.p. Ketamine/Xylazine only for the induction at a dose of 75 mg/kg ketamine and 7.5 mg/kg xylazine. For anesthesia maintenance, a solution of Urethane 20% diluted in Saline was injected i.p. at a dose of between 0.3 and 0.8 ml to effect.

### Surgery: Craniotomy

The surgical protocol for this experiment was adapted from previous experiments carried out on binaurally CI implanted rats (Buck et al. 2021; Rosskothen-Kuhl et al. 2021). We shaved the rat’s head and used depilatory cream to clean the skin on the cranium. The stimulus patterns used and analyzed for this study are monaural, and the data described below all consist of responses to stimulation of the contralateral ear. The rats were stabilized in a stereotaxic frame with earbars, a rostral to caudal midline scalp incision was made, and connective tissue and the temporal muscle were removed to expose the temporal bone. Craniotomies were made extending from 2 to 8 mm posterior from bregma, and from 0 to 5 mm ventral and lateral from the squamous suture. This exposed a large portion of the temporal cortex of the rat, including the entire auditory cortex (AC). A head-mount was then fixed to the parietal cranium. This consisted of three 1.2 mm diameter retaining screws that were fixed against a larger M2 screw with dental acrylic. The larger screw served as a head post that could be fixed to a 3D printed head holder.

### Surgery: Cochlear Implantation

With the head fixed as described above, the CI surgery was performed using an approach through the ear canal. The eardrum and the ossicles were removed with fine-tip forceps. To better expose the cochlea, rongeurs were used to remove some of the bone of the inferior wall of the ear canal. A cochleostomy was performed in the center of the cochlea using a 0.6 mm drill bit, targeting the 14 kHz region located in the middle turn of the rat’s cochlea. Through the cochleostomy we inserted a rodent multichannel intracochlear electrode kindly provided by MED-EL, inserting 4 electrode contacts towards the apex. The contacts were spaced in 0.75 mm intervals, and after insertion they were positioned at distances of approximately [4.8, 5.6, 6.4, 7.2] mm from the base of the cochlea (Burda et al. 1988). These points correspond to approximate tonotopic positions from 14 to 5.7 kHz with respect to the rat cochlear tonotopy identified by Mueller M. (1991). The sites were therefore spaced at approximately half octave steps, an appropriate spacing for facilitating comparison to human data (Kan et al. 2013). The basal-most of the four channels served as ground electrode, while the other three served as active sites for the delivery of intracochlear stimulation patterns.

Two tests were then performed to check the proper functioning of the intracochlear electrodes. First, electrode impedances were measured for each of the stimulating electrodes, and impedances below 1 kΩ or above70 kΩ were interpreted as indicative of technical failure such as a possible short circuit or a contact failure, respectively. Second, Electrical Auditory Brainstem Responses (EABRs) were recorded to individual biphasic pulses with a duty cycle of 40 μs anodic, 40 μs zero, 40 μs cathodic. Thresholds of 4 dB or less relative to 100 μA were taken as indicating a functioning electrode.

### Calcium indicator injection

To facilitate subsequent injection of calcium indicator via micropipette we first applied a cotton tip soaked in purified collagenase in a concentration of 50 mg/ml (Sigma Chemical CO., St. Louis, MO) to both craniotomies to soften the dura overlying the AC. Similar collagenase pretreatment had been used in a previous study by Zhu et al. (2002). The cotton tips were left on the dura for approximately 30 minutes, during which time the calcium indicator was prepared following a protocol described in previous studies (Stosiek et al. 2003; Garaschuk et al. 2006; Grienberger and Konnerth 2012). Briefly 50 μg of the calcium-sensitive dye Oregon Green™ BAPTA-1 (OGB-488) was dissolved in 4 μL of a mixture of 80% DMSO and 20% Pluronic F-127 acid. This was then diluted in 36 μL of artificial cerebrospinal fluid and loaded into a micropipette pulled from a glass capillary (Superior Marienfield, Lauda-Königshofen, Germany) on a Narishige PC-100 micropipette puller to a tip width of between 1 and 5 μm.Once the collagenase had been allowed to take effect and the cotton buds had been removed, the OGB solution was pressure injected into 10 to 20 sites spaced across the whole AC at a pressure of 10 psi and at a depth of 0.4 to 0.8 mm from the surface of the dura (Grienberger and Konnerth 2012). The injection sites were chosen so as to achieve a fairly even spread across the craniotomy while avoiding blood vessels and maintaining a minimum distance of approximately 0.5 mm between neighboring sites. During the injections, the level of Calcium Indicator in the pipette was monitored under a microscope to ensure that the volume delivered at each site was typically between 1 and 2, and never more than 5 μL. After the last injection we allowed a period of approximately 30 minutes for the Ca++ indicator dye to diffuse and be taken up by the tissue before data acquisition commenced.

### *In-Vivo* Wide Field Calcium Imaging

The animals were placed in a dark, sound proof box. A custom designed fluorescence microscope (Thorlabs, New Jersey, US) with a 0.5✕ camera tube (WFA4102, Thorlabs) and a 2✕ apochromatic microscope objective (TL2✕-SAP, 0.1 NA, 56.3 mm WD, Thorlabs) was positioned above the craniotomy. The calcium indicator dye was excited by a green LED light source (M490L4, 490 nm, 205 mW Mounted LED, Thorlabs) through a Fluorescein isothiocyanate dichroic mirror (MD499, FITC Dichroic Filter, Thorlabs; reflection band: 470-490 nm, transmission band: 508-675 nm). The fluorescent light emitted by the rat cortex was collected by the objective, passed through a Green Fluorescent Protein emission filter (MF525-39, GFP Emission Filter, Thorlabs; center wavelength: 525 nm), and captured by a passively cooled sCMOS camera (CS2100M-USB, Quantalux 2.1 MP Monochrome sCMOS Camera, Thorlabs) operating at a frame rate of 30 fps in low noise mode. The camera has a native resolution of 1920 x 1080 pixels, but only 1000 x 1000 pixels were used, covering a field-of-view of 5.04 mm^2^. The LED light source and camera were controlled via TTL digital signals sent from the digital IO port of a TDT RZ6 Multi-I/O Processor (Tucker-Davis Technologies, Alachua, US). The same RZ6 also controlled the electrical stimulator used to deliver stimulus patterns to the CI, and it in turn was controlled via an ActiveX interface and custom written Python scripts running on a Windows desktop computer.

### CI Stimuli

The CI stimuli were delivered from a TDT IZ2MH programmable constant current source stimulator (Tucker-Davis Technologies, Alachua, US). For this study, the stimuli consisted of four different three-channel stimulation patterns illustrated in Figure 1. The patterns were chosen to imitate different spectro-temporal patterns resembling simplified representations of different spoken words generated by a clinical device using a CIS-like strategy. In each of the four 1 s long patterns, the combination of active channels changes every 200 ms (each of the 4 “words’ comprises four 200 ms long “phonemes’’). Electrodograms illustrating the four patterns used are shown in Fig 1A. The patterns were delivered at two different pulse rates (300 Hz and 1800 Hz, see Fig 1B), and with two different stimulation modes: Simultaneous Sampling (SS) or Continuous Interleaved Sampling (CIS, see Fig 1C). The aim of the experiment was to determine how easily the cortical responses to the four different stimulus patterns could be discriminated, and to assess the influence of pulse rate and stimulation mode on stimulus discriminability. Each of the four stimulus patterns was presented 15 times at each mode and pulse rate. The different stimulus patterns, pulse rates and modes were presented in a randomly interleaved order, with a 5 s onset-to-onset inter-stimulus interval.

**Figure 1:**
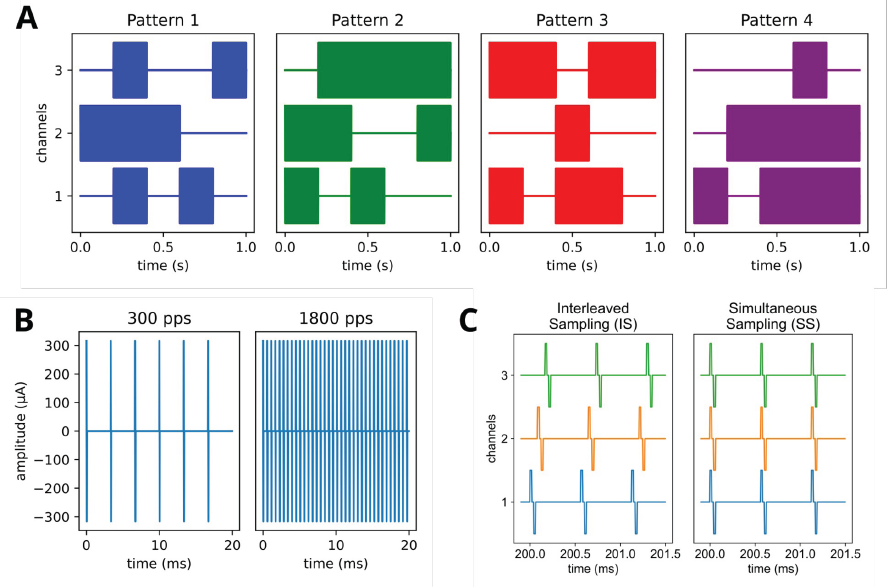
(A) Electrodograms of the four spectro-temporal patterns used in the experiment. (B) Biphasic pulse trains at the two pulse rates used in this experiment (300 pps and 1800 pps). (C) Electrodogram zooming in on a portion of Pattern 1 where all three channels were active, showing the two sampling modes used in this experiment: in interleaved sampling, pulses in neighboring channels are offset in time so only one channel is active at any one time. In contrast, in simultaneous sampling, pulses on neighboring channels are triggered at the same time.

### Data pre-processing and analysis

All recorded optical cortical data were pre-processed and analyzed using custom written Python code. For preprocessing, we first applied a spatial low-pass filter, convolving each collected image with a 25 by 25 pixel, *σ=*5 Gaussian kernel. Then, the filtered images were spatially downsampled by a factor of 5 (OpenCV resize) to reduce the image data size to 100 by 100 pixels. Furthermore, to reduce motion artifacts attributable to the animals’ breathing and heartbeat, the downsampled movies were stabilized by tracking and removing movement relative to the first frame (OpenCV functions calcOpticalFlowPyrLK, estimateAffinePartial2D and warpAffine). Edge artifacts resulting from this image stabilization were removed by trimming 2 rows or columns of pixels from the upper, lower, left and right edges of each frame. After downsampling and image stabilization each 5.04 mm^2^ frame was thus rendered with 96 by 96 pixels.

We then normalized local changes in image brightness by applying the widely used fluorescence ratio change (ΔF/F) calculation to the time series for each pixel (Ren and Komiyama 2021). We first computed the time-varying baseline fluorescence *F_0_(t)* for each pixel as the 5^th^ percentile value over the period from *t-*10 s prior to *t+*10 s. Note that the averaging over 20 s worth of frames is equivalent to a low pass filtering, and consequently *F_0_* can only change very slowly. We found it to be computationally more efficient and just as accurate to compute *F_0_* directly by computing the 5^th^ percentile over image frames in 2.5 s steps and estimate the *F_0_* values between successive 5 s steps by cubic spline interpolation. The *ΔF/F* value for each time *t* is then defined as *(F(t)-F_0_(t))/F_0_(t)* where *F(t)* is the brightness of the pixel at time *t*. In Figure 2A and B we illustrate the conversion of raw luminance time series data to *ΔF/F* for two pixels selected to differ greatly in baseline value and dynamic range. Such differences are more likely to reflect differences in dye loading and illumination than physiological differences in Ca++ dynamics, and the purpose of the *ΔF/F* calculation is to normalize such differences.

**Figure 2:**
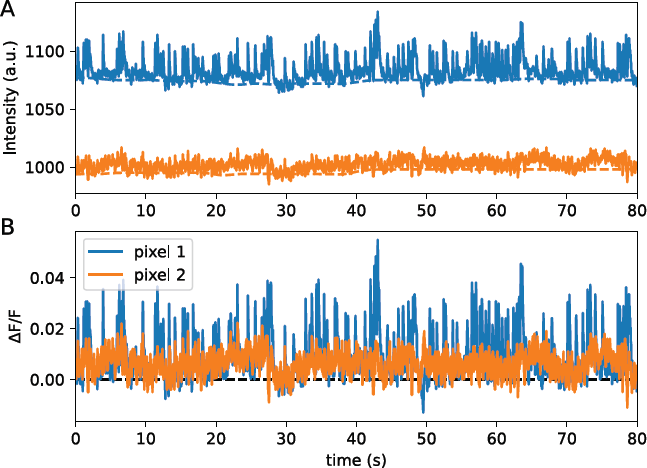
(A) Examples of time series of raw fluorescence signals (solid lines) and their corresponding baseline fluorescence (broken lines) for two discrete pixels (blue and orange lines) for 20 s recording. Each pixel has its own baseline intensity level and dynamic range. (B) Fluorescence ratio (*ΔF/F*) computed for the raw data shown in A. The time series data are baseline corrected and scaled relative to their own baseline.

It is well known that, in sensory cortices, neighboring neurons may share synaptic inputs from incoming axonal arborizations, and therefore often exhibit similar tuning to stimulus parameters. Consequently, neural activity in neighboring pixels recorded with optical techniques tends to be highly correlated. We can take advantage of this correlation structure by performing a principal component analysis (PCA) on the preprocessed data, allowing us to further reduce the dimensionality of the dataset by identifying the dominant patterns of covariance of activity across the cortical surface and re-expressing the 9216 (96 by 96) time series of *ΔF/F* values as only a handful of time series of spatial principal components of covariation. To compute the PCA, each image frame was reshaped into an 9216 element column vector, and these image vectors were stacked horizontally to form a pixel-by-time matrix *M*. A single dataset comprised in the order of 50,000 time frames, and computing the principal components of very large matrices directly can make unfeasibly big demands on computer memory and processing power. We therefore used the IncrementalPCA algorithm implemented in the python library sklearn.decomposition to compute the 15 most important principal components for each image time series.

To assess the discriminability of cortical responses to the different stimulus patterns described above we then performed a classification of single-trial response time series. For each stimulus presentation, we extracted a 60 frame ✕ 15 principal components (PCs) responses matrix, extending for 2 s from stimulus onset, and subjected these response matrices to a simple mean-based classifier. Let *R_i, j_* denote the response matrix recorded in response to the *i*-th presentation of the *j*-th pattern ( *i* ∊ [1, 2, … 15], *j* ∊ [1, 2, … 4]), and let *<u>R</u> _j_* denote the mean response to the *j*-th stimulus pattern, averaged over all responses except *R_i, j_*. Note that leaving the response to be classified out of the computation of group means is important to avoid biasing the classification. Considering each response *R_i, j_* in turn, we then computed the euclidean distance between *R_i, j_* and *<u>R</u> _j_* for each *j,* and simply assigned *R_i, j_* to the stimulus class for which this euclidean distance was smallest. In this manner we constructed 4✕4 confusion matrices, in which the number in each cell simply counts the number of times that a response to the stimulus pattern given by the row of the confusion matrix was assigned to (or “decoded” as belonging to) the stimulus pattern given by the column of the matrix. Perfect classification of the responses will result in confusion matrices with values of 15 in the cells on the main diagonal (given that each stimulus was presented 15 times), and cells on the off-diagonal will be zero. Poorer classification performance will manifest in increased numbers of trials counted in the off-diagonal cells, and a corresponding decline in the counts along the main diagonal. Classification performance can therefore also be summarized as a single percent correct score, computed from the proportion of the total number of responses that were correctly assigned to their places along the main diagonal of the confusion matrix.

## Results

Using the methods just described we collected optical Ca++ responses from the AC of six rats (five right AC recordings, one left AC). As expected, different stimulus patterns did, on average, produce distinct spatio-temporal response patterns across the cortical surface, as illustrated in an example from one of the six rats shown in Figure 3. The example clearly shows that the two different stimulus patterns can evoke, on average, different cortical response patterns. In the example we also note that the responses to the stimuli in SS mode tended to be stronger and more widespread than those to the CIS stimuli, making the differences in the responses to the two patterns more salient in SS than CIS mode. Thus, changes in stimulus parameters such as stimulation mode or pulse rate may well impact the discriminability of cortical responses to different stimulation patterns, and at least in this example, SS appears to result in more easily discriminable responses.

**Figure 3:**
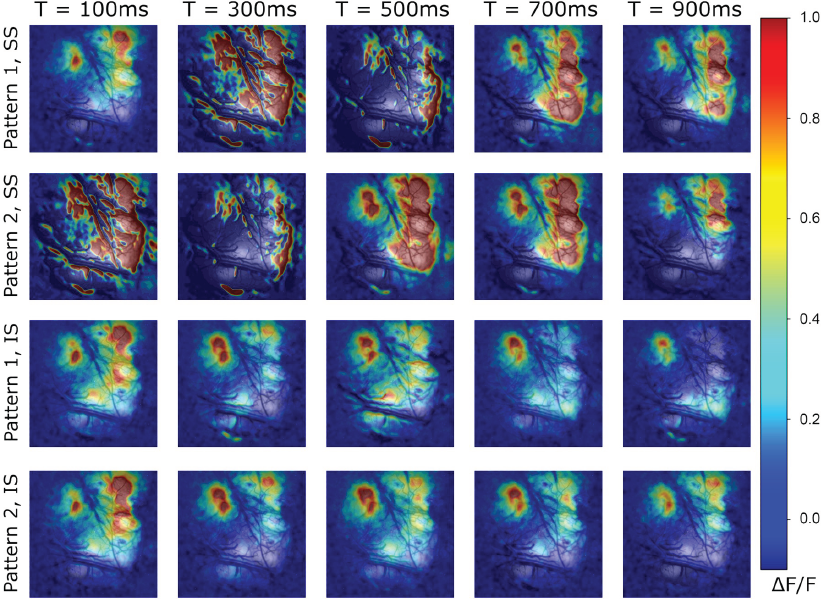
False color maps showing example mean *ΔF/F* responses from one AC superimposed on an image of the AC itself. Responses to stimulus patterns 1 and 2 are shown, presented at 300 pps either in SS or CIS mode, as indicated to the left of each row. The responses are shown at five different time points post stimulus onset, as indicated above each column. The images show 5.04 mm^2^ of cortical area.

To make the qualitative observations made in Fig.3 amenable to quantitative and statistical analysis, the optical data from each animal were first subjected to dimensionality reduction via PCA and then fed into a classification algorithm as described above. The results of the PCA are illustrated in Figure 4. In Fig 4A we plot the percentage of the variance explained by the first 15 principal components (PCs) in the data from each of the 6 animals in our dataset. Our choice to limit our analysis to 15 PCs was somewhat arbitrary, but as can be seen in Fig 4A, the first 15 PCs account for more than 70% of all the variance in the images for 5 out of 6 animals, and in all cases the amount of additional variance explained by more components starts to asymptote by between 4 and 7 PCs in all cases, suggesting that 15 PCs are likely adequate to capture the bulk of the spatial pattern information inherent in our data.

**Figure 4:**
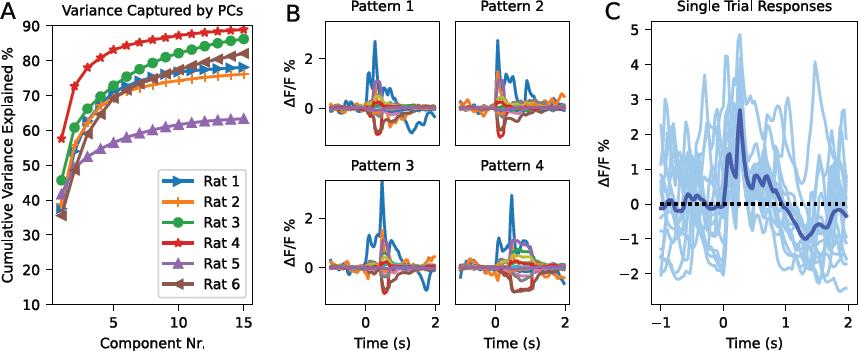
Principal component analysis of the optical responses. A: Cumulative variance explained by the first 15 principal components for each of the 6 rat ACs in our dataset. B: Averaged principal component responses from one example AC to each of the four stimulus patterns presented at 300 pps in SS mode. The responses of each of the 15 principal components are shown superimposed. C: Single trial responses for first principal component for the case shown in the top left of B (stimulus pattern 1 at 300 pps in SS mode). The light blue lines show responses in individual trials, the dark blue shows the mean response over all 15 trials.

Fig 4B shows example responses of the first 15 PCs to the four different stimulus patterns, presented at 300 pps in SS mode, averaged over the 15 presentations of each of these stimuli, for a single AC. The mean responses to the different stimulus patterns exhibit relatively obvious differences, suggesting that the stimuli ought to be identifiable by their respective, distinctive cortical response patterns. However, we need to bear in mind that, in real life situations, auditory brains cannot rely on averaged responses to many presentations to identify a particular sound, but instead have to attempt an identification based on single trial responses, and these are noisy, as is illustrated in the example in Fig 4C. Nevertheless, decoding the stimulus identity from single trial PC responses is often possible using the simple mean-based classifier algorithm described above. Figure 5 illustrates this, showing confusion matrix examples for each of the 6 rat cortices for the 1800 pps stimuli presented in SS mode. Note that the results are fairly consistent across all 6 animals. Classification was perfect for Rat 1 and near perfect for Rats 2 and 5, and although it was slightly worse for Rats 3, 4 and 6, classification was nevertheless well over 60% correct in all cases, very much better than the 25% chance performance one might expect if the classification failed completely.

**Figure 5:**
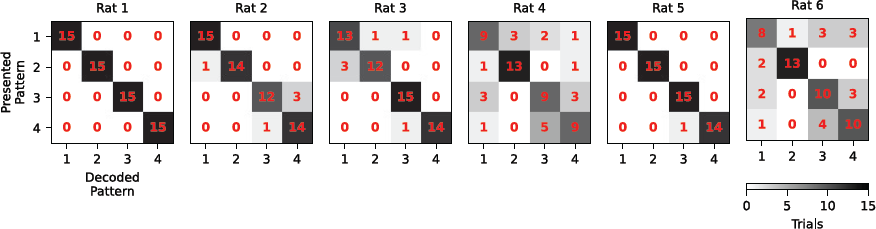
Examples of confusion matrices for each of the 6 animals for the 1800 pps SS stimulation case. The numbers in red give the number of times (out of a maximum of 15) that a response to the stimulus pattern indicated by the row number was identified as belonging to the stimulus indicated by the column number. The grayscale indicates the same number for ease of visualization.

Having shown in Fig 5 that single-trial response classification can give consistent results across our cohort of 6 animals, we can now ask whether the chance of correctly identifying a stimulus pattern depends on pulse rate or stimulation mode. In Fig 6 we show confusion matrices, pooled across all 6 animals, for each of the four stimulation parameters (two pulse rates by two modes) tested, and we can readily observe that the probability of correct classification was substantially higher when the stimuli were presented in SS rather than CIS mode. In contrast, changing pulse rates from 300 to 1800 pps had no very obvious effect: then mean correct classification results for SS were 87.2% and 85.8% for 300 and 1800 pps respectively, while the corresponding values for CIS were 41.1% and 49.4%.

**Figure 6:**
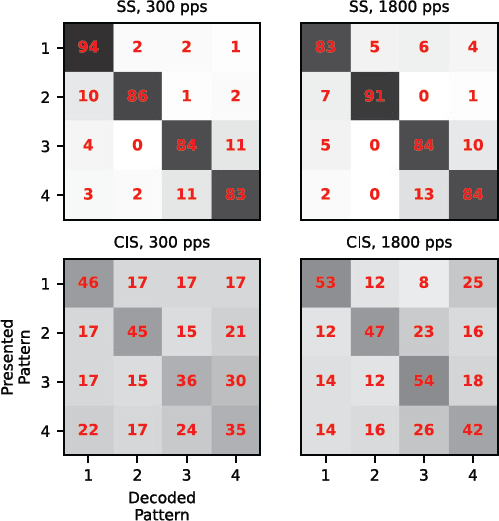
Confusion matrices, as in Fig 5 but pooled over all animals, for each of the four different stimulation parameters tested (top row: SS mode, bottom row CIS mode, left column 300 pps, right column 1800 pps). The numbers in red give the percentage of times (out of a possible maximum of 90) that a response to the stimulus pattern indicated by the row number was identified as belonging to the stimulus indicated by the column number. The grayscale indicates the same number for ease of visualization.

In Figure 7 we summarize the classification performance as a percent correct score for each of the 6 animals and each of the 4 stimulation parameters. It is readily apparent that all 6 animals exhibit the same trends, namely notably higher classification scores when stimuli were presented in SS rather than CIS mode, but essentially similar scores for 300 or 1800 pps. We fitted a linear mixed-effects model for the classification performance with mode (SS vs. CIS), pps (300 or 1800), and their interaction as fixed factors, and with random intercepts for the rats. There was a highly significant main effect of mode (F(1,20)=90.7, p=7.2*10^-9^). The main effect of pps was not significant (F(1,20)=0.082, p=0.78), nor was the interaction between mode and pps (F(1,20)=2.0, p=0.17).

**Figure 7:**
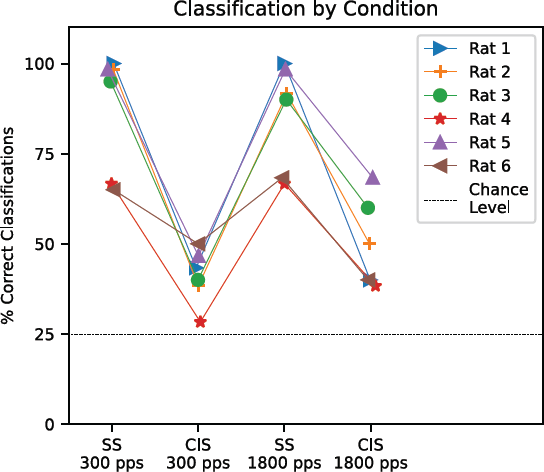
Classification performance (percentage of trials counted along the main diagonal of confusion matrices) computed separately for each of the stimulation parameters as indicated on the x-axis and for each of the 6 animals as indicated on the legend. The thin dotted black line indicates the expected chance performance level of 25% for comparison.

## Discussion

### Discriminability of Ca++ widefield responses as a metric of CI encoding

The main objective of this study was to assess the potential of cortical Ca++ imaging to evaluate different CI stimulation strategies, based on the assumption that differences in cortical response patterns should correlate with how easily the stimuli evoking the response patterns can be discriminated perceptually. The assumption appears reasonable in the light of previous electrophysiological work linking differences in cortical responses to psychophysical performance (Engineer C. et al. 2008; Walker et al. 2008; Shetake J. et al. 2011; Centanni et al. 2014). The results just presented demonstrate that the approach is also highly feasible, within a set of limitations and caveats which we will discuss below. In Fig 3 we suggested that different CI stimulus patterns can evoke clearly distinct patterns of cortical Ca++ widefield responses, in Figs 4 and 5 we argued that these widefield responses can be neatly summarized in a modest number of PC time series that lend themselves for evaluation by classifier algorithms, and in Figs 6 and 7 we saw that the outcomes of such a classification can reveal clear, systematic dependencies of the discriminability of cortical responses on parameters of the stimulation strategy used when encoding the stimuli. Together, these observations suggest that widefield Ca++ imaging may be a very useful, and so far entirely under-utilized tool for assessing certain aspects of CI stimulation strategies.

One important advantage of this approach is that it can sample a large amount of cortical tissue, collecting potentially a large amount of information. Another is that the physiological measurements are completely unaffected by electrical artifacts, even when highly complex, multichannel stimulation patterns are used. Algorithms for removing electrical artifacts from electrophysiological recordings under CI stimulation have steadily improved (O’Shea and Shenoy 2018; Najafabadi M. et al. 2020), but no method is 100% effective, and many of the methods currently in use work well only under a limited set of circumstances, such as a given range of pulse rates or constant pulse rates or very precise synchronization between stimulation and recording hardware. Complementary studies using Ca++ imaging can therefore be helpful in allaying any residual doubts whether the artifact removal in any particular key study was indeed “good enough”, particularly if the studies used complex multichannel stimulation patterns. Ca++ widefield imaging is also a comparatively inexpensive and easy method that can be readily performed on experimental animals with a background of hearing experience or prior experience of electrical intracochlear stimulation that the experimenter can precisely control.

Calcium imaging also has limitations compared with ECoG and Electrophysiology methods. One significant shortcoming when using Ca++ indicator dyes like OGB-488, which we used here, is that these dyes are prone to fairly rapid photo bleaching, which limits the total recording time to a maximum of around 90 to 120 minutes, depending on the strength of the illumination. This poses a significant limitation on the amount of time available for data collection. Another shortcoming of the technique relative to electrophysiological measures is the relatively slow timecourse of the Ca++ dependent signals, which precludes analyses on millisecond timescales.

### Effects of pulse rate and stimulation mode

In the experiment described here, we encoded 4 different stimulus patterns using two different pulse rates (300 and 1800 pps) and two different modes, SS and CIS. We found that changing pulse rate produced no statistically significant effect, and that is probably unsurprising given that previous psychophysical studies reported very similar speech perception thresholds for a variety of different pulse rates (Shannon et al. 2011). In contrast, changing stimulation mode from CIS to SS made a large and statistically highly significant difference, in a direction that may at first seem surprising: in our experiments, SS produced much more easily discriminable cortical responses than CIS. This finding may appear to strongly contradict well established clinical practice, given that at present, essentially all clinically available devices depend largely or entirely on coding strategies that are effectively variants of the CIS algorithm (Carlyon R. and Goehring 2021).

While we believe that our observation that SS produced much more easily discriminable responses is correct in the specific case examined here, we do not claim that this result would generalize to a wider set of circumstances. By comparing CIS and SS head-to-head in an experiment where all other factors are held equal we merely illustrate that whether temporally interleaving channels is advantageous or not depends on context, and to our knowledge there are no compelling experiments demonstrating the superiority of CIS over any other CI sound coding strategy which does not impose a ban on at least occasional simultaneous pulses at multiple electrodes. The original paper by Wilson et al. (1991) describing the CIS strategy merely states “the CIS strategy addresses the problem of channel interactions through the use of interleaved non-simultaneous stimuli’’ but offers no explanation or references indicating how important or effective it is to “address channel interactions” in this manner.

Efforts to understand, and minimize, channel interactions in cochlear implants go back a long way, with an influential paper by Shannon (1983) being one of the earliest to demonstrate the potential for channel cross-talk with loudness and masking interactions, and numerous other papers since have used human psychoacoustic or physiological measures to investigate channel interactions (e.g (Chatterjee and Shannon 1998; McKay et al. 2001; Middlebrooks 2004; Bierer 2007; Tang et al. 2011)). One of the few papers to attempt a direct comparison of SS and CIS stimulation is a study by (De Balthasar et al. 2003), which did indeed demonstrate that channel interactions can be larger when pulses are simultaneous rather than interleaved. However, how detrimental even quite large channel interactions are is not always obvious. A study by Goehring et al. (2021) is instructive in this context: these authors used a “blurring” of input channels to investigate how increasing channel interactions would affect speech perception in CI users. Increasing channel interactions through blurring did indeed negatively affect speech discrimination, but, interestingly, only when the blurring was quite pronounced, spreading information from any one CI channel evenly over 5 or 6 adjacent intracochlear electrodes. Blurring information over 3 or 4 adjacent channels had no significant effect, which indicates that deviations from channel independence have to become quite substantial before they really substantially impair the effectiveness of multichannel cochlear implants.

Another important factor to bear in mind is that electrical crosstalk or physiological interactions between channels are not the only types of interactions that will reduce the independence of CI channels. For CI channels to be truly independent information channels, each channel would have to be able to deliver its own independent temporal pattern code. The highest channel capacity for a channel delivering pulses is achieved if any channel can deliver a pulse of any amplitude at any time. However, CIS solves the issue of pulse collision avoidance in neighboring channels by forcing all channels in the array to conform to the same temporal grid of regular, fixed pulse intervals, sacrificing the possibility to encode temporal fine structure features of incoming sounds with sub-milisecond accuracy. Recent developments in CI research strongly suggest that leveraging the full potential of precisely timed pulses could result in clinically relevant improvements in binaural hearing (Rosskothen-Kuhl et al. 2021; Buck A. et al. 2023) and periodicity pitch perception (Goldsworthy and Shannon 2014; Goldsworthy and Bissmeyer 2023) for CI users, but to deliver such advantages in a multichannel implant capable of effective speech encoding may require lifting the restriction that each channel must operate at the same, fixed, underlying pulse rate regardless of the nature of the input signal, even if this makes it much harder to avoid occasional simultaneous pulses in more than one channel.

The fact that SS performed better than CIS in the case we examined here can most likely be explained by the fact that SS produced a summation of electric fields, which resulted in a stronger stimulation of the auditory pathway (compare top and bottom rows in Fig 3) which in turn leads to a better signal-to-noise ratio in the neural representation of the stimulation patterns. In our case, the advantage of a more effective stimulation at equal pulse amplitudes appears to have more than compensated for any detrimental effects by the expected increase in channel interaction. So while we do not doubt that the enhanced channel interactions provoked by SS may well be disadvantageous in some circumstances, our result nicely illustrates that there can also be circumstances in which relaxing the demand that there must be no simultaneous pulses on multiple channels can be advantageous.

## Conclusion

All CI coding strategy designs involve compromises and trade-offs. The current state-of-the-art in clinical processor design strongly favors trying to optimize spatial separation of channels over leveraging the auditory pathway’s exquisite sensitivity temporal patterns. While the limitations of current approaches in poor pitch perception and binaural hearing are well documented, developing radically new approaches is hard. Trials on human volunteers are confounded by the fact that these volunteers will have thoroughly adapted to their specific clinical devices and may have become desensitized to stimulation features that their devices do not provide. Animal electrophysiology can be revealing as long as electrical artifact removal does not become too difficult, but coding strategies with multiple channels each running independently at variable rates would create electrical artifacts of daunting complexity. Here we have shown that Ca++ imaging of cortical neural responses can provide a valuable additional research tool for evaluating and comparing the effectiveness of different CI coding strategies in the absence of clinical trial confounds or electrical artifacts.

## Acknowledgements

This work was supported by the Hong Kong Health and Medical Research Fund (HMRF grant 07181406), the Shenzhen Science Technology and Innovation Committee (JCYJ20180307124024360), as well as the Martin Lee Centre for Innovations in Hearing Health at Macquarie University .We are grateful to Ms Kamini Sehrawat for very helpful advice on the experimental preparation. IN was partially supported by ISF grant 1126/18, and is the Milton and Brindell Chair in Brain Sciences at Hebrew University.

